# The impact of parvalbumin interneurons on visual discrimination depends on strength and timing of activation and task difficulty

**DOI:** 10.1101/2024.06.07.597911

**Authors:** Lilia Kukovska, Jasper Poort

## Abstract

Parvalbumin-expressing (PV) cells are the most common class of inhibitory interneurons in the visual cortex. They are densely connected to excitatory cells and play important roles in balancing cortical circuit activity and learning. PV cell activation is a tool to inactivate cortical regions to establish their role in visual processing. However, it is not established how moderate activation affects behaviour and how effects depend on activation strength, timing and task difficulty. We therefore investigated how these three major factors affect performance of mice in a go/no-go orientation discrimination task. We tested discrimination performance with different strength and timing of PV cell activation in V1 and with different task difficulty levels. We found that PV cell activation improved performance in easy discriminations when stimulating with moderate laser powers only during the initial 120 ms from stimulus onset, corresponding to the initial feedforward processing sweep across the cortical hierarchy. In the same animals, PV cell activation did not aid performance in difficult discriminations. However, in both easy and difficult discriminations, optimal behavioural performance required undisturbed late phase activity beyond 120 ms, highlighting the importance of sustained activity in V1. Combining the optogenetic activation of PV cells with two-photon imaging showed that behavioural changes were associated with changes in stimulus response selectivity in V1. Thus, our results demonstrate that early and sustained activity in V1 is crucial for perceptual discrimination and delineate specific conditions when PV cell activation shapes neuronal selectivity to improve behaviour.

- Effects of moderate optogenetic PV cell activation on behaviour are time window specific
- Benefits of increased PV cell-mediated inhibition depend on task difficulty
- Optimal behavioural performance requires sustained V1 activity in both easy and difficult discriminations
- Behavioural changes with PV cell activation are reflected by changes in the selectivity of V1 neurons

## INTRODUCTION

GABAergic inhibitory interneurons shape the activity of excitatory pyramidal (Pyr) cells and are critical for learning and balanced activity in the cortical circuits (Froemke, 2015; Isaacson and Scanziani, 2011). Altered inhibition is associated with a range of perceptual and behavioural impairments (del Pino et al., 2018). However, the mechanisms by which cortical inhibition modulates perception and behaviour are not well understood.

The primary visual cortex (V1) contains neurons highly selective for visual features such as orientation (Hubel and Wiesel, 1959; Niell and Stryker, 2008), and is required for visual discrimination (Glickfeld et al., 2013; Poort et al., 2015). GABAergic inhibition in V1 can increase feature selectivity (Katzner et al., 2011; Li et al., 2008; Nelson et al., 1994; Sillito, 1979, 1975), and elevated cortical inhibition facilitates discrimination learning (Ishikawa and Saito, 1978).

It is not yet established which inhibitory cell type is driving these processes. The mouse cortex contains three major types of GABAergic interneurons (Xu et al., 2010), expressing parvalbumin (PV), somatostatin (SOM), and vasoactive intestinal peptide (VIP). Accounting for ∼40%, PV cells are the most common interneuron type and most densely connected to local Pyr cells (Pfeffer et al., 2013; Tremblay et al., 2016). By innervating Pyr cells at the soma and axon initial segment, PV cells play a role in gain control and feedforward inhibition (Markram et al., 2004; Tremblay et al., 2016). PV cells are therefore strategically positioned to facilitate neuronal processing.

Disrupting the balance of inhibition can also be detrimental to sensory perception. Stronger inhibition from PV cells silences V1 to abolish behavioural performance (Glickfeld et al., 2013; Jin et al., 2019; Poort et al., 2015). Importantly, the effect of V1 silencing can depend on the timing of visual processing and the difficulty of the task (Javadzadeh and Hofer, 2022; Resulaj et al., 2018). While silencing the earliest V1 activity abolishes perceptual discrimination, delaying the silencing by 80 ms after the visually-evoked response onset allows discrimination above chance (Resulaj et al., 2018). Difficult discriminations require longer delay than easy discriminations (Resulaj et al., 2018). These findings suggest that distinct roles are played by the early feedforward and later feedback phases of cortical processing to enable visual discriminations (Lamme and Roelfsema, 2000). In contrast to strong inhibition, moderate (non-silencing) inhibition can enhance feature selectivity. A previous study suggested that PV cell activation increased orientation selectivity (Lee et al., 2012),. However, other studies found that PV cells influence the response gain of Pyr cells without altering their orientation selectivity (Atallah et al., 2012; El-Boustani and Sur, 2014; Ingram et al., 2019; Wilson et al., 2012). More recent work suggests that PV-Pyr cell interactions in V1 are plastic and change with visual discrimination learning, providing the local network with more selective inhibition after learning (Khan et al., 2018).

To establish the role of PV cells in visual discrimination and their involvement in the different phases of visual processing, we experimentally manipulated three major parameters in the same group of animals. We trained head-fixed mice on a visual discrimination task and used optogenetics to precisely manipulate the activity of PV cells in V1 during task performance. We systematically varied the strength and timing of optogenetic stimulation, and trained animals to perform both easy and difficult visual discriminations.

We discovered that moderate levels of PV cell activation in V1 enhanced performance dependent on the strength and temporal window of activation. We did not observe discrimination improvements when PV cell activation lasted throughout the entire stimulus presentation. Despite testing a wide range of stimulation levels, performance was always impaired, showing a gradual reduction in behavioural responses to the rewarded orientation. Delaying optogenetic stimulation relative to stimulus onset by up to 180 ms, to target only the late secondary V1 response, greatly minimised this reduction but caused a significant increase in responses to the unrewarded orientation. While performance was above chance level, consistent with a previous study (Resulaj et al., 2018), we found that it was nevertheless impaired. This highlights the importance of sustained undisrupted late phase V1 processing that is modulated by recurrent and feedback input. In contrast, limiting PV cell activation to the early phase of V1 processing (the initial 120 ms of stimulus presentation) caused a significant improvement in discrimination with a reduction in behavioural responses to the unrewarded orientation. Increased task difficulty elevated the incorrect responses to both stimuli, thus worsening the overall performance, and these effects were amplified by PV cell activation. Interestingly, there was a strong interaction of the effects of PV cell activation with task difficulty. We found no strength or timing of activation that improved performance during difficult discriminations.

Finally, to compare the local V1 network activity during the entire stimulus and early phase photostimulation conditions, we repeated the experiments with simultaneous two-photon calcium imaging from neurons in layer 2/3. Consistent with our behavioural findings, we discovered that moderate PV cell activation in easy discriminations decreased stimulus selectivity when it lasted throughout the entire stimulus condition and increased selectivity when activation was limited to the first 120 ms. In difficult discriminations, changes in selectivity did not clearly correlate with changes in behaviour.

Overall, our findings show that the impact of PV cells on visual discrimination and V1 response selectivity critically relies on the strength and timing of activation and task difficulty.

## RESULTS

To investigate the effect of varying the strength and timing of PV cell activation and task difficulty on visual perception, we trained mice to perform a visual go/no-go discrimination task. Head-fixed mice were trained to run on a cylindrical Styrofoam treadmill (Figure 1A) and initiate trials by maintaining running speed above a threshold for 0.5 – 0.9 seconds. Each trial consisted of a grey background inter-trial interval (3 6 s), followed by a two-second presentation of one of two orientations separated by 90 degrees (Figure 1B). Mice were rewarded for licking a spout during the ‘go’ stimulus. There was no punishment for licking to the ‘no-go’ stimulus, except during training when stimulus presentation was extended by up to 6 s during a timeout period. Trial outcomes (Figure 1B) were used to calculate behavioural performance (*d’*, see Methods).

**Figure 1.**
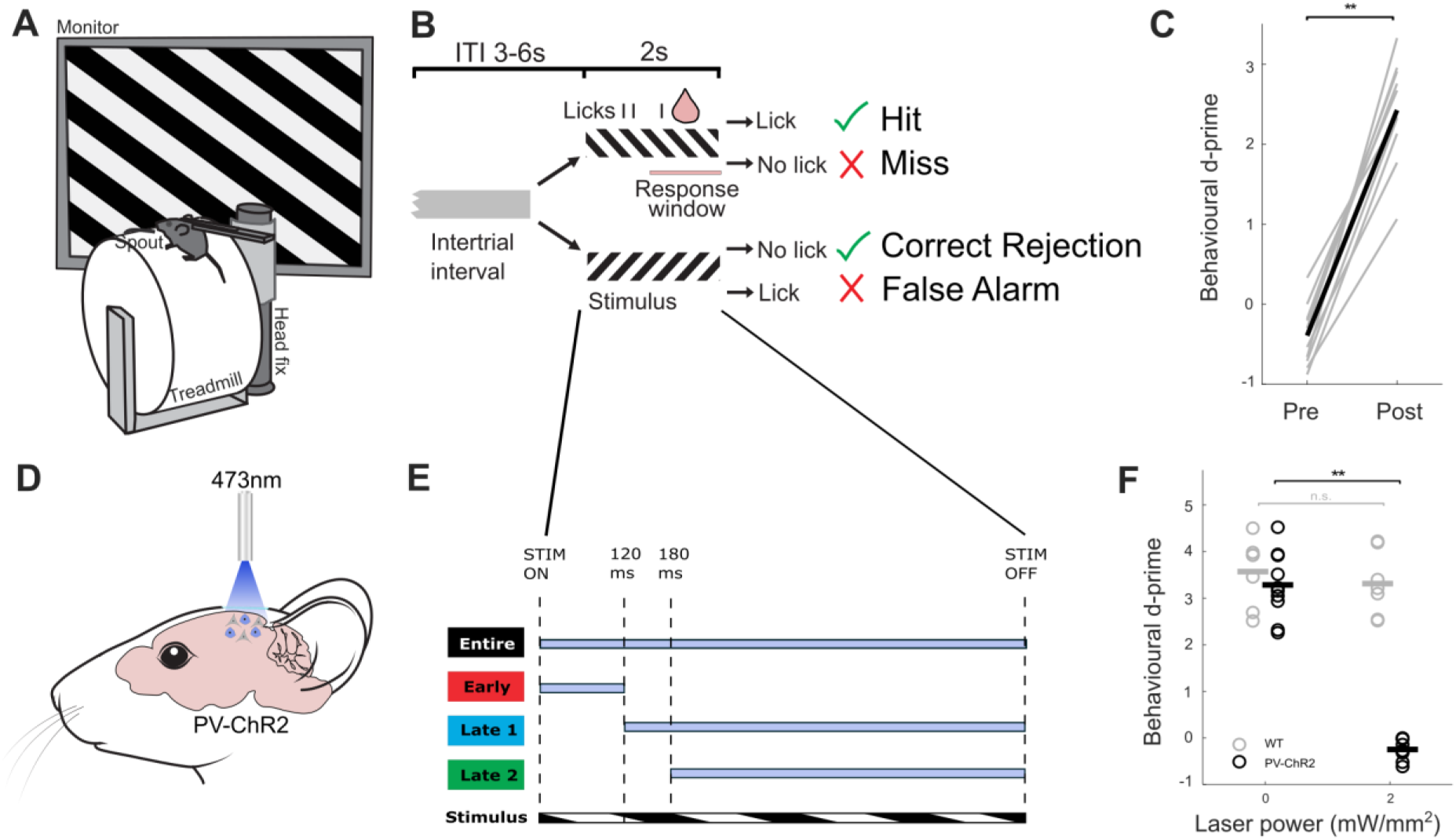
Learning of a visual discrimination task and optogenetic manipulations in head-fixed mice. A) Schematic of setup. Mice are head-fixed on a cylindrical Styrofoam treadmill and static gratings are presented on a monitor to the right eye (contralateral to photostimulated side). A spout located in front of the mouse delivers milk during rewarded trials. B) Schematic of visual discrimination task. Static grating (2 s) is presented following a variable grey background inter-trial interval (ITI) lasting (3 – 6 s). For ‘go’ trials, the reward is delivered when licks are detected during the response window (1 2 s after stimulus onset). Depending on the presence or absence of licks, ‘go’ trials are classified as hit or miss and ‘no-go’ trials as false alarms and correct rejections, respectively. C) Pre- and post-learning behavioural discrimination performance, *d’*, averaged across mice (Wilcoxon signed-rank test, p = 0.002, N = 10). Grey lines represent individual mice. D) Schematic of optogenetic method. Optic fibre is centred over V1 to deliver blue light (473 nm) to activate Channelrhodopsin-2 (ChR2) expressed in all parvalbumin-positive interneurons of transgenic mice (PV-Cre::Ai32). E) Schematic of stimulation protocol. Optogenetic stimulation is delivered relative to stimulus onset (STIM ON) and offset (STIM OFF). ‘Entire’ condition (black) lasts during the entire stimulus presentation; ‘early’ condition (red) lasts from stimulus onset to 120 ms; ‘late1’ condition (blue) lasts from 120 ms to stimulus offset; ‘late2’ condition (green) lasts from 180 ms to stimulus offset. Each condition was tested in separate sessions (interleaving different power levels). F) Discrimination performance, *d’*, in wild-type mice (WT, N = 6, 7 sessions) and transgenic mice (PV-ChR2, N = 10, 10 sessions) at no laser stimulation (0 mW/mm^2^) and silencing laser power (2 mW/mm^2^). Circles, individual sessions; line, mean.

Mice initially licked indiscriminately during the grey background inter-trial interval and stimulus presentation (Figure S1A, top panels), resulting in high proportions of hit and false-alarm trials (Figure S1C). As training progressed, mice gradually learned to withhold licking to non-rewarded stimuli (Figure S1A, middle panels). Eventually mice learned to exclusively lick in response to the ‘go’ stimulus (Figure S1A, bottom panels). Running speed also changed stereotypically over the course of training (Figure S1A, right panels) and mice learned to slow down upon stimulus presentation and accelerate for unrewarded stimuli. Successful discrimination was evident by the significant increase in behavioural *d’* across mice (Figure S1B). Mice performed at chance level in their first training session (-0.39 ± 0.12) and reached an average performance of *d’* > 2 after training (2.43 ± 0.21) (Figure 1C, Wilcoxon signed-rank test, p = 0.002, N = 10).

### Strong activation of PV cells in V1 impairs discrimination performance

Strong activation of PV cells is frequently used as a tool to silence cortical activity and study the functional role of brain areas (Cone et al., 2019; Glickfeld et al., 2013; Javadzadeh and Hofer, 2022; Kaplan et al., 2016). However, different strengths of PV cell activation in V1 may differentially affect behaviour (Lee et al., 2012; Poort et al., 2015). To examine how manipulating PV cell activity over a wide range of laser strengths affects perception, we used photostimulation of PV cells expressing Channelrhodopsin-2 (ChR2) in transgenic mice. We estimated the boundaries of V1 (see Methods) and sealed all adjacent areas with black paint. We then centred an optic fibre above the craniotomy to deliver blue 473 nm light to V1 (Figure 1D). To exclude the possibility of behavioural effects arising due to non-specific effects from the light itself (Owen et al., 2019), we used silencing laser powers in mice lacking ChR2 as a control (Figure 1F). At 2 mW/mm^2^ laser power illumination, discrimination in control wild type mice remained high (*d’* 3.57 ± 0.28 to 3.31 ± 0.26, Wilcoxon signed-rank test, p = 0.297, N = 6, 7 sessions). Moreover, the laser light did not affect pupil size dynamics (Figure S1D). In contrast, discrimination in transgenic PV-ChR2 mice was fully abolished (Figure 1F; *d’* 3.01 ± 0.16 to -0.25 ± 0.07, Wilcoxon signed-rank test, p = 0.002, N = 10, 10 sessions).

Optogenetically manipulating V1 activity could potentially induce behavioural responses by itself. To determine whether mice responded to PV cell activation alone, we activated PV cells at different intensity levels in trained mice during grey screen without visual stimulus. PV cell activation alone did not induce licking behaviour (Figure S1E).

### PV cell activation during entire stimulus presentation impairs performance at all non-silencing levels

Since PV cells are well situated to modify feedforward transmission (Rudy et al., 2011), and have been linked to altering response gain and orientation selectivity, and form stimulus-selective ensembles with Pyr neurons (Atallah et al., 2012; Khan et al., 2018; Lee et al., 2012; Wilson et al., 2012), we hypothesised that low levels of activation could sharpen sensory representations to improve discrimination. We therefore tested PV cell activation at a range of non-silencing laser powers (< 0.5 mW/mm^2^, see Methods), while mice performed easy discriminations (Figure 2, 90 degrees angle difference between ‘go’ and ‘no-go’ stimuli). When light was delivered throughout the entire stimulus presentation, visual discrimination progressively decreased as a function of laser power. Performance declined from 2.86 ± 0.14 (*d’*, Figure S2A) to between 1.5 to 2 units of *d’* lower at the four highest powers (Figure 2A, ‘entire’). As expected (Glickfeld et al., 2013; Poort et al., 2015), the strongest PV cell activation at 0.43 mW/mm^2^ reduced behaviour to low performance levels (*d’* 0.91 ± 0.19, Figure S2A), confirming that V1 activity is required for visual orientation discrimination. Interestingly, the effect was mostly mediated by a sharp decrease in licking to the rewarded stimulus (Figure 2B), reaching as low as 0.49 ± 0.08 (hit rate, Figure S2B). Despite a small significant reduction (Figure 2C), licking to the unrewarded stimulus remained relatively stable around 0.22 ± 0.04 (Figure S2C). Notably, individual mice showed variability in false-alarm (FA) rate change across laser powers with some increasing and others reducing their licking rate (Figure S2E), suggesting that mice adopted different strategies when presented with the same sensory limitations.

**Figure 2.**
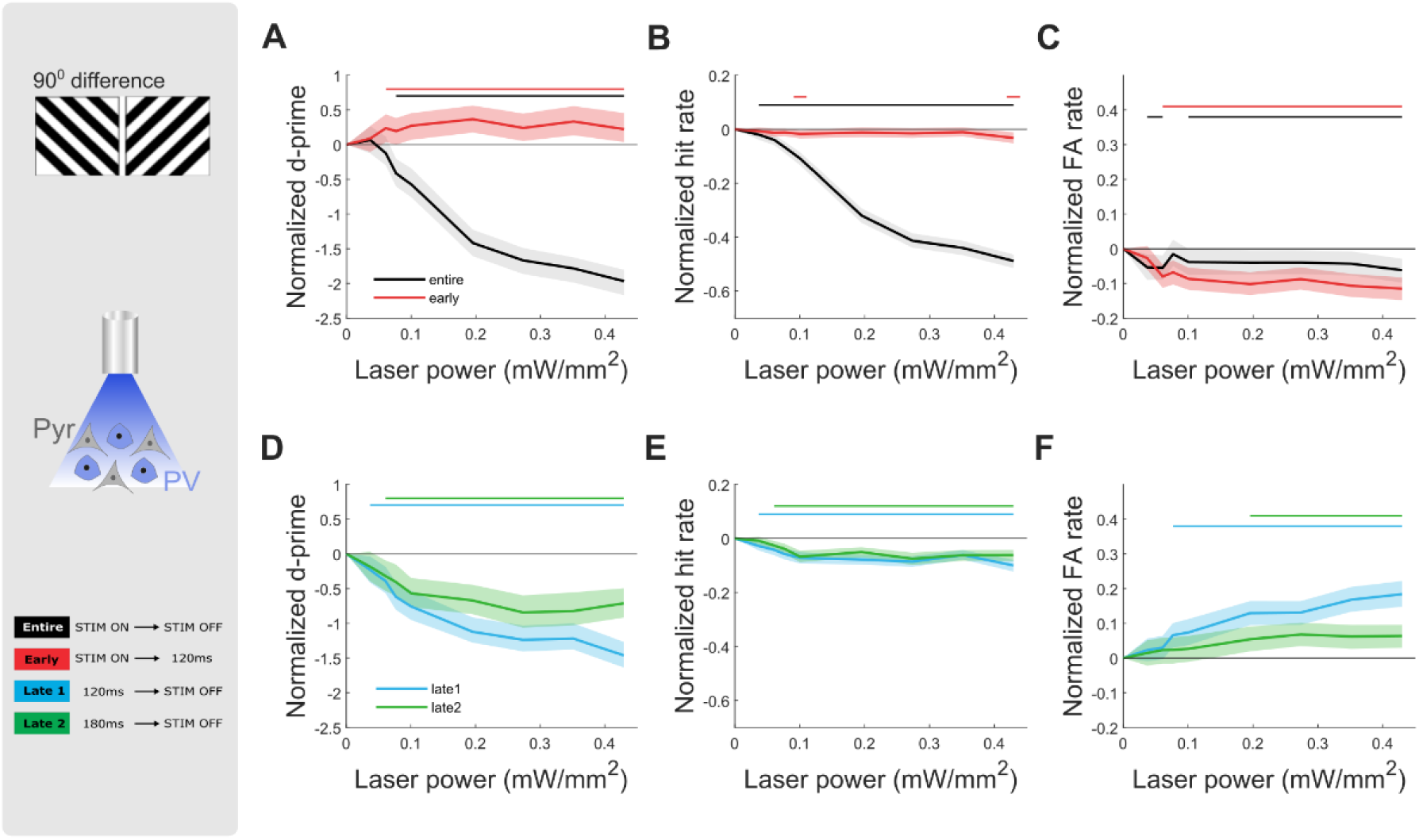
Stimulating PV cells at non-silencing levels during entire stimulus presentation or late phase V1 activity impairs discrimination performance, whereas stimulating during early phase V1 activity improves performance. Grey panel: Example of grating stimuli separated by 90 degrees for easy task (top), schematic of optogenetic manipulations activating only PV cells expressing ChR2 (middle), optogenetic stimulation timings for all four conditions aligned to stimulus onset (STIM ON) or offset (STIM OFF) (bottom). A) Normalized average discrimination performance, *d’,* B) normalized average probability of correct ‘go’ trials (hit rate), and C) normalized average probability of incorrect ‘no-go’ trials (false alarm rate), as a function of laser power for ‘entire’ window (black, N = 10 mice, 41 sessions) and ‘early’ window (red, N = 10 mice, 37 sessions) conditions. Data is normalized by subtracting the value of the 0 power condition (see Methods). D) to F) same as in A) to C) but for ‘late1’ window (blue, N = 10 mice, 40 sessions) and ‘late2’ window (green, N = 8 mice, 32 sessions) conditions. All plots: shaded line is the 95th percentile of data shuffled 1000 times, using bootstrapping with replacement. Horizontal bars with corresponding colours indicate significant deviation (p < 0.05) from no laser stimulation control condition.

We did not observe an activation level of PV cells that significantly enhanced discrimination. Where previous findings suggested that improvements arise from the significant increase in hit rate (Lee et al., 2012), in our task, mice already showed a high hit rate (Figure S2B, above 0.95 for 0 mW/mm^2^). We therefore determined whether improvements in behavioural *d’* were masked by a ‘ceiling effect’. We restricted the analysis to include only sessions with a hit rate lower than 0.95 for the baseline (no laser) condition and confirmed our initial findings with no behavioural improvements following PV cell activation (Figure S2D).

### PV cell activation restricted to the early phase V1 activity improves performance at all non-silencing levels

It has been previously suggested that the earliest visually evoked activity in V1 is sufficient to enable perceptual discriminations (Resulaj et al., 2018). We therefore investigated if moderate PV cell activation restricted to this initial 120 ms following stimulus onset (Figure 2, ‘early’), without interfering with sustained V1 activity that is modulated by recurrent and feedback signals, improved visual discrimination. Indeed, we observed a consistent significant improvement in behavioural *d’* across all non-silencing laser powers (Figure 2A), from 3.11 ± 0.12 up to 3.45 ± 0.16 (Figure S2A). These behavioural improvements were also present across individual mice (Figure S2F, in eight of ten mice), and resulted from a significant decrease in FA rate across all levels (Figure 2C), from 0.20 ± 0.03 down to 0.09 ± 0.02 (Figure S2C), and a stable hit rate (Figure 2B), with values remaining high in the ranges of 0.95 ± 0.02 and above (Figure S2B).

We next wondered whether these perceptual benefits would disappear at high levels of inhibition. Indeed, performance at silencing laser powers was impaired in two out of three mice (Figure S2G) and the third mouse delayed its response for the same duration as the laser stimulation (Figure S2H).

### Undisturbed late phase V1 activity is required for optimal performance

Even at lower hierarchical levels such as V1, visual cortical neurons remain active after participating in the initial feedforward sweep of information following the stimulus onset (Lamme and Roelfsema, 2000). While the early response window is thought to encode basic visual features, activity in the late phase response is more closely associated with the behavioural choice of the animal (Lamme and Roelfsema, 2000). Since we found that moderate photostimulation restricted to the early phase V1 activity improved discrimination, we asked whether increasing PV cell-mediated inhibition during the sustained V1 activity window could also benefit performance, by targeting top-down influences or biasing the behavioural choice. To spare the initial feedforward sweep, we restricted light delivery to specific time windows, delayed by 120 ms or 180 ms from stimulus onset, and lasting until stimulus offset (Figure 1E, ‘late1’ and ‘late2’ respectively). This approach allowed us to simultaneously investigate the effects of varying the strength and timing of PV cell activation.

Irrespective of the delay with which PV cells were activated, we found that performance dropped gradually with increasing laser power (Figure 2D). Behavioural *d’* for late1 condition decreased from 3.36 ± 0.12 in the absence of stimulation to reach values between 2.0 and 2.4 at the strongest stimulations (Figure S2A). On the other hand, *d’* for late2 condition decreased less, from similar levels of 3.40 ± 0.13 to levels between 2.7 and 2.9 (Figure S2A). This divergence in performance was most evident at laser powers over 0.2 mW/mm^2^. In both delayed conditions the hit rate was maintained at high levels (although slightly reduced, Figure 2E) over 0.90 ± 0.03 across all tested laser powers (Figure S2B), contrasting the entire window condition where hit rate was reduced. The reduced *d’* was mainly explained by a change in FA rate (Figure 2F). The FA rate in the late1 condition gradually increased over laser powers to double from 0.16 ± 0.03 at baseline to 0.33 ± 0.05 at 0.43 mW/mm^2^ (Figure S2C). The FA rate in the late2 condition was maintained at a low rate across all laser powers at around 0.17 ± 0.04 (Figure S2C). Interestingly, all mice either sustained or increased their FA rate (in contrast to the entire window condition where mice adopted variable strategies, Figure S2E).

The performance of mice was less impaired when they were given more time to perceive the stimulus without manipulating the activity of V1. A previous study has shown that the first visually evoked spikes in V1 are sufficient for making perceptual decisions, resulting in performance above chance levels (Resulaj et al., 2018). Importantly, our results demonstrate that easy discriminations rely on unaltered late phase V1 activity to achieve maximum performance.

### Difficult discriminations do not benefit from PV cell activation

After establishing the effects of varying the strength and timing of PV cell activation on easy discriminations, we tested difficult discriminations where stimuli were separated by 15 degrees. Mice showed significantly reduced performance but were still able to discriminate above chance. We trained mice on the new stimuli until they achieved *d’* > 1.5 (average performance during testing *d’* > 2, Figure S3A). We found that the effects of the entire (Figure 3A-C) and both delayed (Figure 3D-F) window conditions were similar to those reported in easy discriminations (Figure 2), with reductions in *d’* of a comparable size.

**Figure 3.**
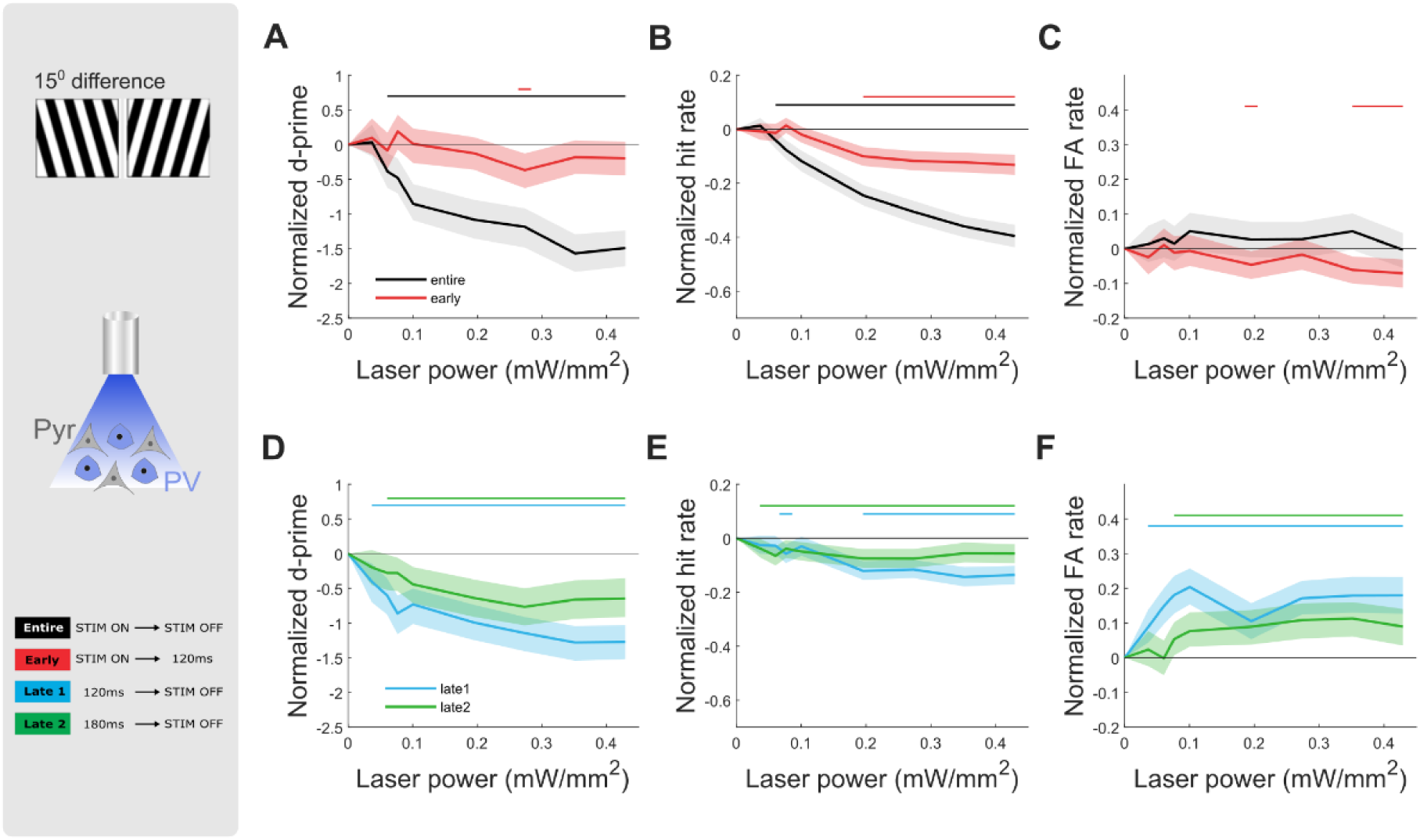
Stimulating PV cells at non-silencing levels does not improve performance of difficult visual discriminations. Grey panel: Example of grating stimuli separated by 15 degrees for difficult task (top), schematic of optogenetic manipulations activating only PV cells expressing ChR2 (middle), optogenetic stimulation timings for all four conditions aligned to stimulus onset (STIM ON) or offset (STIM OFF) (bottom). A) Normalized average discrimination performance, *d’*, B) normalized average rate of correct ‘go’ trials (hit rate), and C) normalized average rate of incorrect ‘no-go’ trials (false alarm rate), as a function of laser power for ‘entire’ window (black, N = 7, 24 sessions) and ‘early’ window (red, N = 8, 28 sessions) conditions. D) to F) same as in A) to C) but for ‘late1’ window (blue, N = 7, 25 sessions) and ‘late2’ window (green, N = 7, 25 sessions) conditions. All plots: shaded line is the 95th percentile of data shuffled 1000 times, using bootstrapping with replacement. Horizontal bars with corresponding colours indicate significant deviation (p < 0.05) from no laser stimulation.

After restricting laser stimulation to the early window, we saw no improvements in *d’* across laser powers (Figure 3A). The hit rate significantly decreased with stronger PV cell activation at powers greater than 0.2 mW/mm^2^ (Figure 3B), counteracting the beneficial effects of a slightly decreased FA rate (Figure 3C).

Since difficult discriminations require sustained V1 activity (Resulaj et al., 2018), we wondered whether longer PV cell stimulation would be beneficial. To test this, we introduced two new windows lasting from stimulus onset to either 180 ms or 340 ms. We found that the changes in *d’*, hit and FA rates were comparable to the standard early window (Figure S3D-F). Thus, we did not identify manipulations in the strength or timing of PV cell activation that improved performance in difficult discriminations.

To further understand the effects of moderate PV cell activation in V1 on behaviour, we investigated the influence on various aspects of the decision-making (Figure 4). First, we investigated whether improvements in discrimination could be explained by a speed-accuracy trade-off strategy (Heitz, 2014; Rinberg et al., 2006). Second, we investigated how the laser illumination influenced the animal’s decision criterion (Stanislaw and Todorov, 1999). Third, we investigated the effect of PV cell activation on the profile of licking, running and pupil size. Fourth, we investigated the effect of PV cell activation on different parameters of a simple model describing decision-making dynamics.

**Figure 4.**
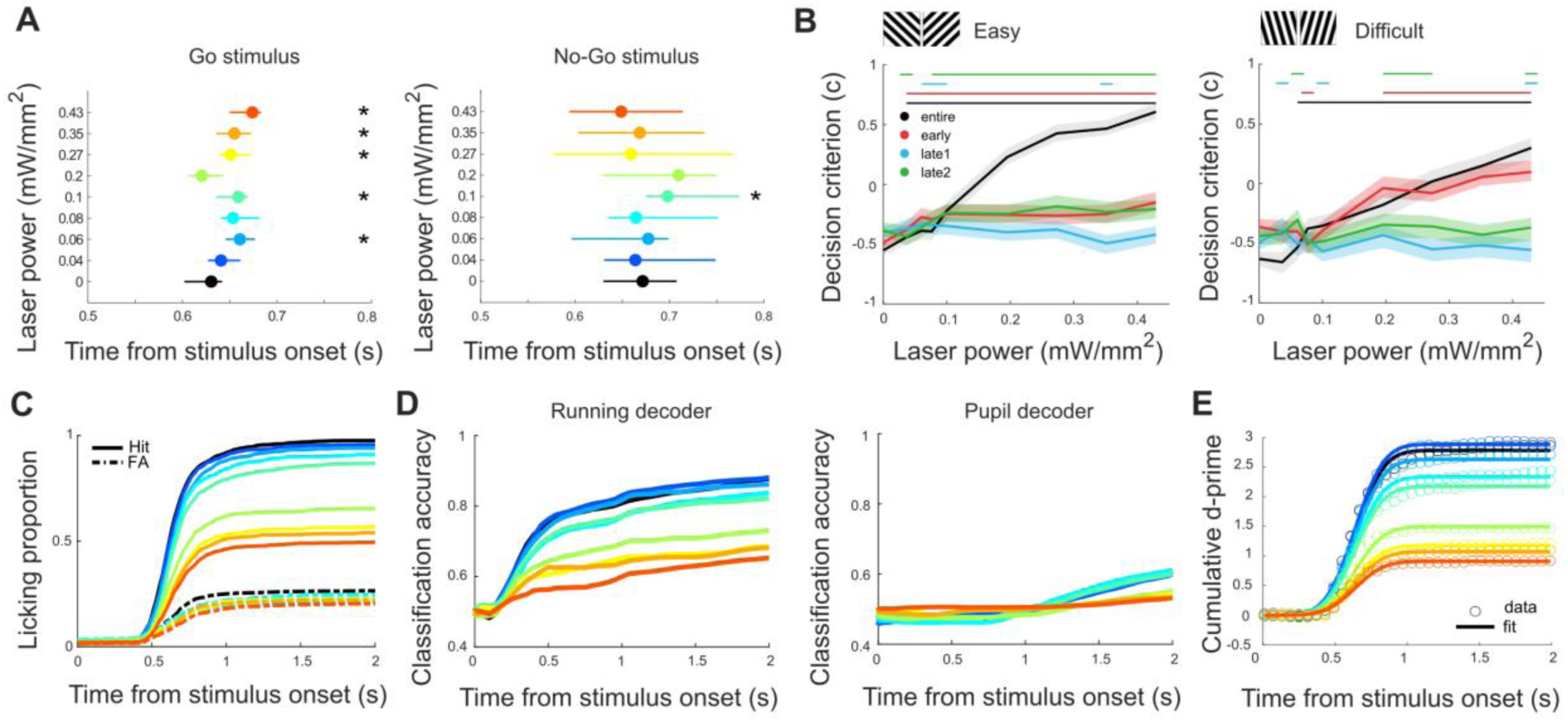
PV cell activation effects on different decision-making components. A) Response latency for time to first lick in laser off (0 mW/mm^2^) versus laser on (0.04 – 0.43 mW/mm^2^) in the early window condition for ‘go’ (left panel) and ‘no-go’ (right panel) stimuli. Circle, median; line, 95% CI of bootstrapping. Asterisks on right of axis indicate significant deviation (p < 0.05) from no laser stimulation control condition. B) Decision criterion *c* across time windows in easy (left panel) and difficult (right panel) discriminations. Shaded line is the 95% CI of data shuffled 1000 times, using bootstrapping with replacement. Horizontal bars with corresponding colours indicate significant deviation (p < 0.05) from no laser stimulation. C) Licking proportion in response to the presentation of the ‘go’ (solid line) and ‘no-go’ (dashed line) stimuli. Plots in C) to E) show data for the entire window condition, with colours ranging from blue to red corresponding to increasing laser powers as indicated in A). D) Time course of classification accuracy of running speed (left panel) and pupil size (right panel) linear decoders (probability of correctly identifying ‘go’ versus ‘no-go’ trials), based on the cumulative evidence of running speed or pupil diameter for different PV cell activation levels. E) Normal cumulative distribution function (solid line) with fixed mean and standard deviation as a model fit for cumulative *d’* (circles) across stimulus presentation (for visualisation purposes, every second data point is shown). The number of mice and sessions is the same as in Figures 2 and 3.

### Behavioural improvements are not due to speed-accuracy trade-off strategy

We asked whether the improvements from early window PV cell activation could result from mice adopting a speed-accuracy trade-off strategy, with slower responses reflecting more accumulated evidence and improved performance (Heitz, 2014; Rinberg et al., 2006). To quantify reaction times for each trial, we measured lick latencies following stimulus onset (Figure 4A). Although some PV cell activation levels showed changes in the median time to first lick to the rewarded stimulus, delays were minor (20 to 40 ms, Figure 4A, left panel). Moreover, there were no significant delays in the responses to the unrewarded stimulus (Figure 4A, right panel), despite discrimination improvements arising from reductions in FA rate (Figure 2C). Thus, a speed-accuracy trade-off strategy did not explain the results.

### PV cell activation affects decision criterion

In signal detection theory (Stanislaw and Todorov, 1999), the sensory process can be characterised by a sensitivity parameter, *d’*, and the decision process by a decision criterion parameter, *c* (see Methods). While *d’* quantifies the separation of two distributions, the decision criterion *c* quantifies whether the animal is more liberal (negative criterion, animal more likely to respond) or more conservative (positive criterion, animal less likely to respond). We investigated whether the changes we observed in *d’* were associated with changes in the decision criterion. Regardless of task difficulty, the criterion in the entire window condition shifted from negative to positive values, i.e. lick to non-lick trends (from -0.55 ± 0.08 at 0 mW/mm^2^ to 0.61 ± 0.22 at 0.43 mW/mm^2^ in the easy task and from -0.63 ± 0.10 at 0 mW/mm^2^ to 0.30 ± 0.34 at 0.43 mW/mm^2^ in the difficult task; Figure 4B, black). Thus, with the increased strength of PV cell activation, mice became more conservative in responding. Positive shifts in criterion were also observed in the early window condition but only when mice performed difficult discriminations (from -0.36 ± 0.11 at 0 mW/mm^2^ to 0.10 ± 0.18 at 0.43 mW/mm^2^; Figure 4B, red). Thus, mice adopted a more conservative strategy when faced with more complex discriminations. Interestingly, despite interleaving laser powers within recording sessions, criterion values were different, suggesting mice rapidly adapted their behaviour on a trial-by-trial basis (Aberg and Herzog, 2012). Together, these results show that PV cell activation in V1 not only influences perceptual evidence but also the decision processes.

### Similar PV cell activation effects on running speed and licking responses

We next investigated how PV cell activation in V1 influenced other behavioural readouts such as running speed and pupil size. After learning, running shows a stereotypical speed profile, with an early detection component after stimulus onset – a reduction in running speed for both the ‘go’ and ‘no-go’ stimulus, and a late component discriminating between the ‘go’ and ‘no-go’ stimulus (Figure S1A, right bottom panel). Pupil dynamics have been linked to decision-making performance in other tasks (Cavanagh et al., 2014; Lee and Margolis, 2016). To compare the effects of PV cell activation on running and pupil size with the effects on the licking proportion during both stimulus conditions (Figure 4C) we used a linear decoder (see Methods). The decoder predicted based on the running speed and pupil size the identity of each trial (classifying it as a ‘go’ or ‘no-go’ trial), enabling comparison of classification accuracy and licking proportion. We found that the running speed decoder classified trials with accuracy above 80% in the absence of optogenetic PV cell manipulations (Figure 4D, left panel; here shown for the entire window condition).The profile of running speed resembled the profile of the licking proportions (Figure 4C), with a clear separation between laser conditions suggesting similar effects on both behavioural readouts. Interestingly, changes in running speed diverged earlier than the licking proportion responses. In contrast, in our task, pupil size was a poor predictor of stimulus identity, with decoder performance accuracies below 60%, and was not systematically modulated by the laser manipulations (Figure 4D, right panel).

In conclusion, running speed accurately predicts discrimination performance and is modulated by PV cell activation in V1, providing an early readout of the animal’s intention to respond to the stimulus by licking. In contrast, pupil dynamics are not a good predictor of behavioural performance.

### PV cell activation modulates amplitude of evidence accumulation

We next determined which function described the dynamics of decision-making and the effects of PV cell activation. We quantified the decision-making dynamics with the cumulative *d’* (quantifying the *d’* up to a certain time, see Methods). We found that the normal cumulative distribution function provided good fits with three parameters (the mean describing the onset, the standard deviation quantifying the variability, and the amplitude describing the strength of evidence integration). We created three versions of the model. In model one, the mean, standard deviation, and amplitude were all free to vary across laser powers. In model two, the mean and standard deviation were fixed across laser power conditions. In model three, only the amplitude was fixed. Model two with fixed mean and standard deviation but allowing changes in the amplitude (Figure 4E, here shown for the entire window condition), significantly outperformed the other two models across all time windows and difficulty levels (median R^2^ of model 2 = 0.995; 2.5^th^ and 97.5^th^ percentiles of model 1 = 0.984 and 0.987; 2.5^th^ and 97.5^th^ percentiles of model 3 = 0.705 and 0.748). In conclusion, the evidence accumulation was well described by a simple function. Interestingly, the onset and variability of evidence accumulation remained relatively consistent across all conditions, while the amount differed across laser powers.

### Changes in neural selectivity reflect behavioural performance in easy but not difficult discriminations

Since improvements in easy discriminations were not due to mice adopting a speed-accuracy trade-off strategy, we wondered whether changes in performance were associated with changes in the selectivity of the V1 network to stimulus. To simultaneously record the activity of V1 neurons while activating PV cells, we repeated the same experiments in a different cohort of PV-Cre mice, where we expressed a calcium reporter GCaMP7s and a red-shifted opsin ChrimsonR in layer 2/3 of V1 (Figure 5, see Methods). We confirmed that red light (639 nm) illumination increased PV cell responses and suppressed Pyr cell responses to the visual stimulus and the effects scaled with laser power (Figure 5A, example cells).

**Figure 5.**
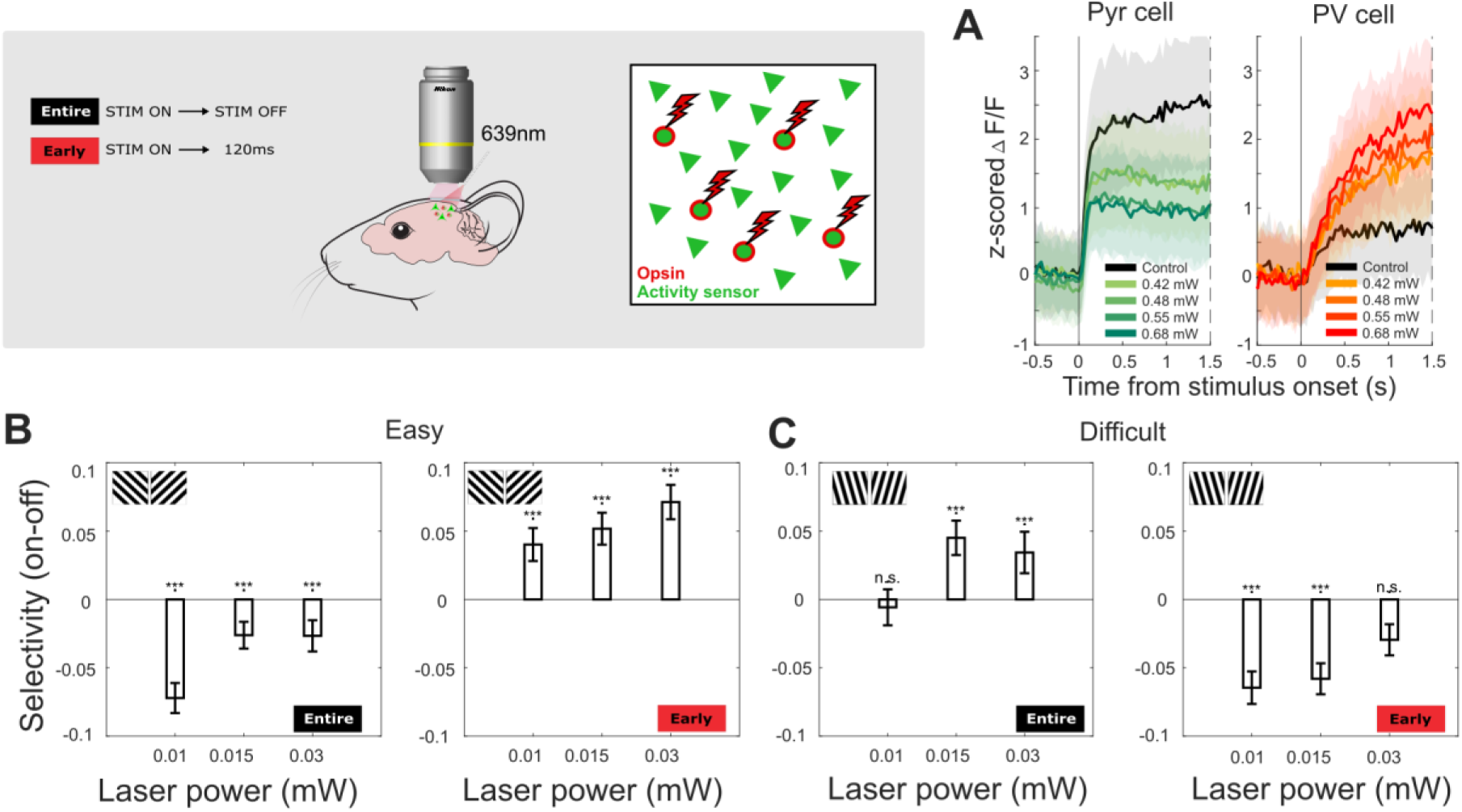
Changes in neural selectivity caused by PV cell activation reflect behavioural performance in easy but not difficult discriminations. Grey panel: Optogenetic stimulation timings for entire and early window conditions aligned to stimulus onset (STIM ON) or offset (STIM OFF) (left), schematic of optogenetic method combined with 2-photon imaging (middle), schematic of laser activating only opsin-expressing PV cells while calcium sensor reports the activity of all neurons (right). A) Average response of two example neurons to optogenetic laser illumination during the entire stimulus duration (0 – 2 s). The activity to the visual stimulus of the identified Pyr cell is suppressed, whereas the PV cell is increased. Shading, SEM. B) Median laser-induced changes in absolute selectivity for entire (left panel, N = 686 cells, 3 mice) and early (right panel, N = 383 cells, 3 mice) time window conditions, for easy task, significant at all laser powers (Wilcoxon signed-rank test, p < 0.002). Error bars, SEM. C) is same as B) showing entire (left panel, N = 337 cells, 2 mice) and early (right panel, N = 313 cells, 2 mice) time window conditions but for difficult task (Wilcoxon signed-rank test, entire 0.01 mW p = 0.986, early 0.03 mW p = 0.060, all other p < 0.002).

Response selectivity in V1 has been previously linked to higher discrimination performance (Poort et al., 2015). The selectivity is defined as the difference in the response to ‘go’ and ‘no-go’ stimulus divided by the pooled standard deviation (see Methods). We therefore investigated the selectivity for each time window and laser power separately. When mice performed easy discriminations, the selectivity across all laser powers decreased when activating PV cells during the entire stimulus presentation (Figure 5B, left panel; largest change for laser power 0.01 mW = -0.07 ± 0.01, p < 0.002, n = 686 cells, 3 mice) and increased when restricting activation to the early window only (Figure 5B, right panel; largest change for laser power 0.03 mW = 0.07 ± 0.01, p < 0.002, n = 383 cells, 3 mice), reflecting the previously reported behavioural changes (Figure 2A). A clear match between reductions in behavioural performance was not observed in difficult discriminations, despite significant changes in absolute selectivity (Figure 5C). Thus, our results show that increased behavioural performance during PV cell activation in the early time window is associated with increased response selectivity in V1.

## DISCUSSION

Cortical inhibition facilitates visual processing and aids visual learning and discrimination in both humans and rodents (Edden et al., 2009; Frangou et al., 2018; Goel et al., 2018; Ishikawa and Saito, 1978). However, how activation of PV cells benefits or perturbs visual discrimination has not been established. Our study is the first to examine in the same animals the combined effects of three major factors that affect discrimination performance. We tested the effects of increasing PV cell activity during different phases of V1 processing on visual discriminations with varying difficulty. We found that non-silencing PV cell activation can benefit behaviour. However, effects were time window and task difficulty specific, and associated with changes in the response selectivity of V1 neurons.

Multiple studies have strongly activated PV cells as a tool to silence V1 and study its involvement in behaviour (Glickfeld et al., 2013; Jin and Glickfeld, 2020). Changing the timing of silencing of V1 has helped to determine the epochs in which V1 contributes to behaviour (Javadzadeh and Hofer, 2022; Resulaj et al., 2018), leading to the idea that the first spikes during the feedforward sweep of information processing are already sufficient to enable perceptual decisions. On the other hand, using non-silencing levels helps reveal the function of inhibitory interneurons in shaping the activity of principal neurons and behaviour (Cone et al., 2019; Lee et al., 2012). However, the functional role of PV cells has been under debate (Atallah et al., 2012; El-Boustani et al., 2014; Lee et al., 2012; Wilson et al., 2012), and how PV cells contribute to the different phases of V1 processing is not known. Here, we mapped the full range of PV cell activation strengths at different time windows of V1 activity. Our results show that moderately increased cortical inhibition following non-silencing PV cell activation could benefit behaviour, but only when it is restricted to the feedforward sweep of V1 processing. The improvements were not a result of mice adopting a speed-accuracy trade-off strategy, which would have likely affected both easy and difficult discriminations (Abraham et al., 2004; Heitz, 2014). Rather, behavioural changes were linked to changes in the selectivity of V1 neurons, suggesting a possible limitation in the role of PV cells when executing tasks of varying complexity. Generally, PV cells are more broadly tuned than Pyr cells, which is thought to be due to their non-specific network connectivity (Hofer et al., 2011; Kerlin et al., 2010; Scholl et al., 2015). Potentially, this makes their tuning precision insufficient to enhance Pyr cell activity during more difficult discriminations.

In our task, mice indicated their decision by licking a reward spout. We also measured the effect of moderate PV cell activation on running speed and pupil size to understand the influence on the decision-making process. It has been shown that studying movements during behaviour provides a continuous readout of the evidence accumulation and the internal decision dynamics (Kane et al., 2023; Pinto et al., 2018). We found that changes in running speed preceded licking, accurately predicted the choice of the animal, and were equally modulated by PV cell activation. Pupil size, previously linked to motivation, decision confidence, and behavioural state (McGinley et al., 2015; Vinck et al., 2015), did not predict stimulus identity in our task. This supports previous work dissociating running from pupil dynamics (Vinck et al., 2015). The mechanisms driving these decision processes can be captured using a simple normal cumulative function. Interestingly, the timing and variability of evidence accumulation appeared preserved despite activating PV cells during different times of visual processing. Instead, the effects were best described by the modulation of evidence strength. Because we simultaneously investigated the impact of three variables in the same cohort of mice, we were able to identify that elevated PV cell-mediated inhibition in V1 did not interfere with the onset of decision-making.

Previous studies have demonstrated an association between the response selectivity of V1 neurons and behavioural performance (Henschke et al., 2020; Poort et al., 2015). We therefore reasoned that the changes in visual discrimination following the moderate activation of PV cells could result from changes in the stimulus response selectivity of the local network. We confirmed our hypothesis that improved discrimination in the early window condition was associated with increases in V1 response selectivity. Similarly, the behavioural impairments in the entire window condition were correlated with decreased selectivity. Interestingly, these effects were only visible in easy discriminations, and we did not find a clear relation between V1 selectivity changes and performance of difficult discriminations. It is possible that complex tasks rely more heavily on higher-level areas to influence decisions, for example on the posterior parietal cortex or the cingulate cortex (Akam et al., 2021; Han and Helmchen, 2024; Licata et al., 2017; Lyamzin and Benucci, 2019; Whitlock, 2017; Zhang et al., 2014).

Traditionally, V1 has been viewed as a region specialized in processing basic visual features such as orientation. However, extensive reciprocal connections between V1 and higher-level visual areas have since been described, suggesting that V1 is a more complex processing area. For example, the activity of V1 neurons can be influenced by behavioural state and running speed (Niell and Stryker, 2010; Saleem et al., 2013; Vinck et al., 2015), spatial navigation (Flossmann and Rochefort, 2021), auditory signals (Bimbard et al., 2023; Chanauria et al., 2019; Williams et al., 2023), predictions (Leinweber et al., 2017) and task context (Hajnal et al., 2023). Although the earliest V1 activity is sufficient for discrimination and enables perceptual decisions above chance (Resulaj et al., 2018), we found that the late phase V1 activity is in fact required for optimal performance. This late phase, i.e. the activity beyond 120ms from stimulus onset, coincides with the time when recurrent and top-down influences from secondary visual and decision-making areas contribute to the local computations to modulate V1 activity (Lamme and Roelfsema, 2000; Wyatte et al., 2014). Feedback to V1 improves visual stimulus encoding required for behaviour (Javadzadeh and Hofer, 2022). Additionally, top-down influences enhance their relative impact with learning (Makino and Komiyama, 2015) and carry predictions that can shape neural responses and facilitate perception (Price et al., 2023; Richter et al., 2023). As a result, activation of PV cells during the late V1 activity phase may perturb such top-down influences and indirectly impair performance.

Our results highlight the important role that PV cells play in shaping neural selectivity and visually guided behaviour. They point to the critical importance of specific time windows of visual processing and calibrated strength of inhibition, and the dependence of behavioural effects on task difficulty. Collectively, these findings underscore the contribution of V1 in both simple and complex visual discriminations and highlight the dynamic nature of cortical inhibition provided by PV cells.

## METHODS

### Animals

All experimental procedures were conducted according to institutional animal welfare guidelines and licensed by the UK Home Office. Mouse lines were obtained from the Jackson Laboratory (PV-Cre, stock number 008069, and Ai32, stock number 012569), and inbred to maintain the original colony or crossed to create a PV-Cre::Ai32 colony (Madisen et al., 2012). Thirteen adult mice underwent surgeries at ages between 7 – 13 weeks. Animals were co-housed in a 12-hour (08:30 – 20:30) reversed light/dark cycle room and provided with cage enrichment and standard diet and water ad libitum. During behavioural testing, mice were food deprived to maintain a minimum of 85% (but typically 90%) of their free-feeding body weight (between 3 – 5 g of standard food pellets per animal per day).

### Surgical procedure

Mice were anaesthetised with 2.5% Isoflurane (1 – 2% for maintenance) with 2 L/min oxygen, administered with pre-operative analgesia of 5 mg/kg injectable Metacam, and positioned in a stereotactic apparatus. A subset of mice also received intraperitoneal injections with a 10 mg/kg dose of Dexamethasone the day before and day of surgery to prevent brain swelling. Body temperature was maintained at 36.5 – 37.0° C throughout the procedure using a heating pad. Ointment (Xailin Night) was applied to the eyes to protect them from drying out. The head was shaved, and a circular piece of scalp was removed to expose the skull which was then cleaned and dried. All mice were fitted with a headpost attached by dental cement (C&B Metabond) to allow head fixation onto the experimental apparatus. A craniotomy was made over the left hemisphere with the centre positioned 2.5 mm left of the midline and the posterior edge aligned with the anterior edge of the transverse sinus. Where viral injections were administered (PV-Cre mice only), Picospritzer (Parker Picospritzer III) was used to inject a total of 800 nL of a 2:18 mixture of AAV-syn-jGCaMP7s-WPRE (Addgene: 104487-AAV1) and AAV-syn-FLEX-rc[ChrimsonR-tdTomato] (Addgene: 62723-AAV5) respectively, in 2 injection sites (coordinates ML2.5 and ML2.3) of 200 nL each at depth of 200 μm and 400 μm from the brain surface. The craniotomy was sealed with a double-layer 4mm-diameter cranial window (two no. 1 thickness windows glued together with index-matched adhesive Norland #71) to provide stable optical access for photostimulation during behaviour (Goldey et al., 2014). A custom-designed 3D-printed cap was used to protect the cranial window. Mice were monitored for five days post-surgery with free access to food and water, with post-operative analgesia of 1.5 mg/ml oral Metacam given in the first three days. After full recovery, mice were food-deprived, and habituated to experimenter handling and head-fixation apparatus, after which training) commenced.

### Behavioural setup

Mice were head-fixed onto a Styrofoam cylinder (Figure 1A). Running was measured with an incremental encoder attached to the cylinder shaft (Kübler). Behavioural task trials were initiated when running exceeded a threshold for a specified time (typically 1 – 2 s). An inter-trial interval (ITI) – grey pre-stimulus illumination, was then presented ranging from 3 – 25 s to discourage timed licking (3 – 6 s for testing). To avoid excessive licking during training, the ITI was extended if mice licked during the grey screen, and up to 6 s timeout was given when mice licked during the unrewarded ‘no-go’ stimulus. A reward spout positioned near the snout of the mouse delivered strawberry-flavoured soy milk via a pinch valve (NResearch), with licks being detected via a piezo disc sensor. Reward was delivered upon first lick only during a reward zone (1 – 2 s after the rewarded ‘go’ stimulus onset) or by an auto-reward trigger at 2 s if mice failed to lick. The stimuli consisted of static black and white gratings with a fixed spatial frequency (0.06 cpd) with 100% contrast and had either 135° or 225° orientation. Stimuli were presented pseudorandomly (no more than four of the same stimulus type in a row). Stimuli were presented with an equal probability of 0.5, except in rare cases where mice failed to show behavioural improvements and the ‘no-go’ stimulus was biased for brief periods with a 0.7 probability to discourage licking. The rewarded stimulus was counter-balanced across mice. Stimuli were presented on a 27” DELL U2715H monitor positioned at 45° to the mouse body axis and 23 cm away from its right eye. Stimulus presentation, reward delivery, and laser stimulation were controlled by a custom-written Matlab script using the Psychophysics Toolbox (Brainard, 1997).

Schematics in Figures 1, 2, 3 and 5 are adapted from SciDraw.io, from the following contributing authors: Emmett Thompson (3925987); Ethan Tyler and Lex Kravitz (3926057); Long Li (5496322); and Manish Kumar (4914800).

### Experimental protocol

For 2 – 3 days, mice were habituated to the cylinder and milk rewards. For 2 – 3 days, mice learned to run on the cylinder (monitor off) uninterrupted and to lick the spout for rewards without visual stimulation. During discrimination training (1 – 3 weeks), mice learned to discriminate between ‘go’ and ‘no-go’ stimuli. Each training session lasted approximately 1 to 1.5 hours (usually between 100 – 400 trials). Mice were tested with optogenetics after achieving *d’* > 2 on 3 consecutive sessions, except for two mice which were tested after 1 such session. This criterion was to prevent overtraining and ensure all four task outcomes (hit, miss, FAs, CRs) were present for analysis. Overtraining will also result in a ‘ceiling effect’, possibly masking behavioural improvements resulting from PV cell activation. Three factors were tested: 1) strength and 2) timing of optogenetic stimulation, and 3) difficulty of discrimination. Each testing session had five interleaved laser powers (including 0 mW/mm^2^ to establish baseline *d’*) and lasted until at least 400 trials were completed. Testing of timing was done in a pseudorandom order.

### Optogenetic stimulation

A Stradus VersaLase laser was used to deliver 471 nm light via a 200 nm core, 0.39 NA optic fibre cable. Laser intensity was calibrated using a standard photodiode power sensor (S121C, Thorlabs). All laser powers are reported as mW/mm^2^. A laser power was first identified to fully silence discrimination (Figure 1F). Lower test powers, including 0.04, 0.06, 0.08, 0.10, 0.20, 0.27, 0.35, and 0.43 mW/mm^2^, were then selected to produce detectable changes in discrimination. The size and boundaries of V1 were estimated using intrinsic imaging data from 12 mice corresponding to previously reported boundaries (Garrett et al., 2014). Black paint was used to seal the window and restrict light delivery to V1. The paint was effective in blocking the laser, with an insignificant leakage of 0.02 µW at the highest testing powers. The optic fibre was centred over V1 at either a 10.7 mm or 17.7 mm distance from the window to deliver uniform illumination. Since the fibre was externally placed, side propagation through the brain to activate neighbouring brain areas or to reach the retina to produce nonspecific effects is unlikely (Yizhar et al., 2011). Additionally, to minimise laser light reaching the eyes, the optic fibre was shielded by a custom-designed 3D-printed cone. Light delivery lasted throughout the entire stimulus duration or for specific time windows (Figure 1E).

### Behavioural and statistical analysis

Task performance was quantified with behavioural *d’* (Stanislaw and Todorov, 1999), with higher values reflecting better performance. Behavioural *d’* and decision criterion *c* were calculated by:

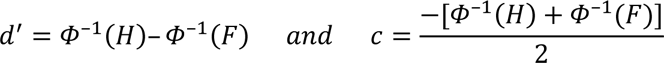

where *Φ* is the normal inverse cumulative distribution function, *H* is the rate of hit trials (licking for rewarded trials during the response window only) and *F* is the rate of false-alarm trials (licking for unrewarded trials during entire stimulus presentation). While *d’* captures perceptual sensitivity, the decision criterion *c* is a measure of the subject’s internal bias to respond; with negative values indicating high bias for ‘yes’ responses and positive values indicating high bias for ‘no’ responses. Lick latencies were determined in trials classified as hit or FA as the first lick detected in the stimulus presentation window (0 – 2 s).

No statistical methods were used to predetermine sample sizes, but the number of mice and sessions included in the behavioural experiment is comparable to previous studies (Danskin et al., 2015; Glickfeld et al., 2013; Poort et al., 2015; Resulaj et al., 2018). All data are presented as mean ± SEM unless otherwise stated. To compare within-subject PV cell activation effects on *d’*, hit, FA, and criterion to the baseline condition (0 mW/mm^2^), we used bootstrap with replacement 1000 times to generate 95% CI (Efron, 1979). Significant effects were defined as non-overlap between the no laser condition and the 95% CI for each laser power.

### Eye tracking

Pupil positions were recorded with a Raspberry Pi camera (model 3B+ or 4), where infrared LEDs illuminated the eye. A small number of eye frames (< 2%) were randomly selected from the whole recording of each session, and pupil extraction thresholds were manually adjusted. The ellipses to track pupil position and diameter were manually verified by inspecting a random subset of samples from each session. See previous studies for more details on eye tracking (Meyer et al., 2020; Poort et al., 2015).

### Decoder analysis

A cumulative decoder was employed to quantify the accuracy with which running speed and pupil size could classify trials as either ‘go’ or ‘no-go’ at time *t* relative to stimulus onset (see Poort et al., 2015). The decoder received inputs of running speed or pupil size aligned to stimulus onset. Thirty trials were taken from each condition to make up the training data, which the decoder then used to construct a model of the running speed or pupil size response to the rewarded and unrewarded stimuli (calculating the mean response µ across trials for each condition). On the remaining trials, the log-likelihood at time *t* that trial *k* belongs to condition *C* (where *C* is the rewarded ‘go’, *R* or unrewarded ‘no-go’ stimulus, *U*) is proportional to:

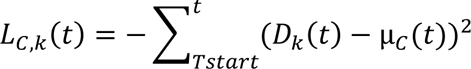

where *D* indicates the running speed or pupil size. The trial was assigned as a rewarded stimulus response if L_R_ > L_U_, otherwise classified as an unrewarded stimulus response. To obtain the cumulative likelihood L_c_ at each time point *t*, the summation only included time points from stimulus onset T_start_ to time *t*. To determine the time point of a detectable divergence, the Wilcoxon rank-sum test was performed on the average speed in individual trials in consecutive nonoverlapping 50 ms windows. The divergence point was defined as the centre of the first window with p < 0.01 followed by at least 4 consecutive windows with p < 0.01.

### Modelling cumulative d-prime

The cumulative d’ was calculated to quantify the *d’* up to a certain timepoint (by calculating the proportion of hits and false alarms at different timepoints relative to stimulus onset). A normal cumulative distribution function (CDF) was used to fit the experimentally observed cumulative *d’* curves (Figure 4E):

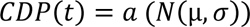

where N is the normal cumulative distribution and the three parameters include a, the amplitude, µ the mean, and σ the standard deviation. Model 1 allowed all three parameters to be varied across laser levels, model 2 only allowed the amplitude to vary (while mean and standard deviation were fixed), and model 3 allowed the mean and standard deviation to be varied (while amplitude was fixed) across laser levels. We used half of the trials for the fitting and the other half of the trials for the testing, using R^2^ to quantify the quality of the fits. The analysis showed that model 2 performed much better than model 3 and slightly better than model 1, indicating that a model with a mean and standard deviation fixed across laser levels, with an amplitude differing across levels, described the data well. Significance was determined by the overlap between the median R^2^ of a model and the 95% confidence interval of the bootstrap distribution of the other models.

### Two-photon calcium imaging

Functional *in vivo* calcium imaging was performed using a two-photon scanning microscope with a 3 mm working distance Nikon 16x 0.8 NA objective and Ti:Sapphire laser (Mai Tai HP Deep See, Spectra-Physics, < 70 fs pulse width, 80 MHz repetition rate) at 920 nm, wavelength for calcium indicator GCaMP7s excitation. A 12 kHz resonant scanner (Cambridge Technology) and an FPGA module (PXIe-7965R FlexRIO, National Instruments) imaged at a 30Hz frame rate to acquire 512 x 512 pixels images covering a 350 x 350 μm field of view. Data was acquired using ScanImage 5.6. A National Instruments DAQ card was used to record triggers for synchronisation of neural responses, stimulus presentation, optogenetic stimulation, eye camera videos, licking, and running. Laser power was set to 15 – 40 mW and was kept consistent throughout consecutive recording sessions. Where identification of tdTomato-tagged PV cells was required, the structural marker tdTomato was excited with a wavelength of 1040 nm.

Imaging commenced no earlier than three weeks after surgery to ensure stable viral expression. A screening session for each mouse was conducted before data collection commenced. During the initial screening session, ThorCam Software (Thorlabs) was used to identify the injection site coordinates guided by blood vessel identification. Retinotopic and orientation mapping helped identify a responsive site for recording with RFs located near the centre of the monitor. A brief recording was made for anatomical identification of tdTomato-labelled PV cells, subsequently used for identifying PV cells. Recordings were made from layer 2/3 neurons at a depth of 150 – 250 μm below the pial surface. Mice were removed from the study if imaging was compromised due to bone regrowth or recording site becoming functionally unresponsive due to overexpression of GCaMP7s, if no single session of *d’* > 1.5 was achieved after extended training, or if neuronal activity was fully silenced even at the lowest possible optogenetic laser power.

### Simultaneous two-photon imaging and optogenetic stimulation

A Stradus VersaLase laser was used to deliver 639 nm light to the imaging site via a 200 nm core, 0.39 NA optic fibre cable coupled with a 20 mm Fiber Optic Cannula. The cannula was positioned at a 30° angle and 7 mm distance from the cranial window and held in place by a micromanipulator. Laser intensity was measured across several levels using a standard photodiode power sensor (S121C, Thorlabs), and calibrated. ChrimsonR-tagged PV cells were stimulated by centering the laser beam over V1 to deliver a uniform illumination. A custom-made 3D-printed cone with attached Blackout Nylon Fabric (Thorlabs BK5) was carefully inserted around the objective and optic cannula to protect the PMTs from external light sources while also eliminating optogenetic laser light from illuminating the eyes.

To allow simultaneous two-photon imaging and optogenetic stimulation within the same neuronal population, the stimulus monitor and optogenetic laser were blanked during the linear phase of the resonant scanner such that the monitor was turned off and the laser was blocked out during the opening of the PMTs shutter, and vice versa.

### Imaging data analysis

Data was pre-processed using Suite2p, where frames were registered to correct for brain motion and active neurons were automatically selected as regions of interest (ROIs) to extract calcium traces for further analysis (Pachitariu et al., 2017). After automated detection, all sessions underwent a manual inspection to ensure identified ROIs only included cell somas and excluded axons, synaptic boutons, or other artefacts. Pixels within each ROI were averaged to obtain raw fluorescence time series *F(t)*. The raw fluorescence values for each ROI were then converted to *Δ F/F*. Neuronal activity was aligned to the onset of stimuli and *Δ F/F* was averaged over 0 – 1 s time window following stimulus onset (excluding the response window 1 – 2 s where reward is delivered) to estimate the response of a cell to oriented gratings. The Wilcoxon signed-rank test was used to determine if the response to the gratings (0 – 1 s) significantly decreased or increased relative to the pre-stimulus baseline activity (-0.5 – 0 s). To calculate the z-scored *Δ F/F* in the 0 – 1 s response window, we subtracted the pre-stimulus response and divided by the pre-stimulus standard deviation. Cells included in Figure 5B/C were cells where the z-scored *Δ F/F* was higher than 0.5.

### Neural selectivity

A selectivity index (SI) was calculated for each cell as (Khan et al., 2018; Poort et al., 2015):

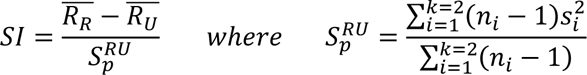

where R_R_ is the averaged stimulus response (0 – 1 s) to the rewarded ‘go’ stimulus and R_U_ is the averaged stimulus response (0 – 1 s) to the unrewarded ‘no-go’ stimulus. The difference is divided by the pooled standard deviation of the responses 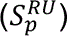, where *n_i_* is the number of trials in i for *k* conditions. A positive or negative value indicates cells with a preference for the rewarded or unrewarded stimulus, respectively. To calculate the median selectivity across cells, we computed the absolute of the selectivity of each cell. For each cell, the absolute selectivity during laser manipulations was subtracted from the absolute selectivity during the control condition (no laser).

## ACKNOWLEDGMENTS

We thank members of the Poort and Beltramo labs for valuable discussions, and John McClure Jr for support throughout the study. We thank Natsumi Homma and Riccardo Beltramo for comments on the manuscript. Schematics were sourced from SciDraw.io and used with permission under the creative commons license (CC-BY). This work was supported by a BBSRC Cambridge DTP targeted studentship (L.K.) and the Wellcome Trust (J.P., 211258/Z/18/Z).

## AUTHOR CONTRIBUTIONS

L.K. and J.P. designed the experiments. L.K. performed the experiments. L.K analysed the data with help from J.P.. L.K. and J.P. wrote the paper.

## SUPPLEMENTARY FIGURES

**Figure S1.**
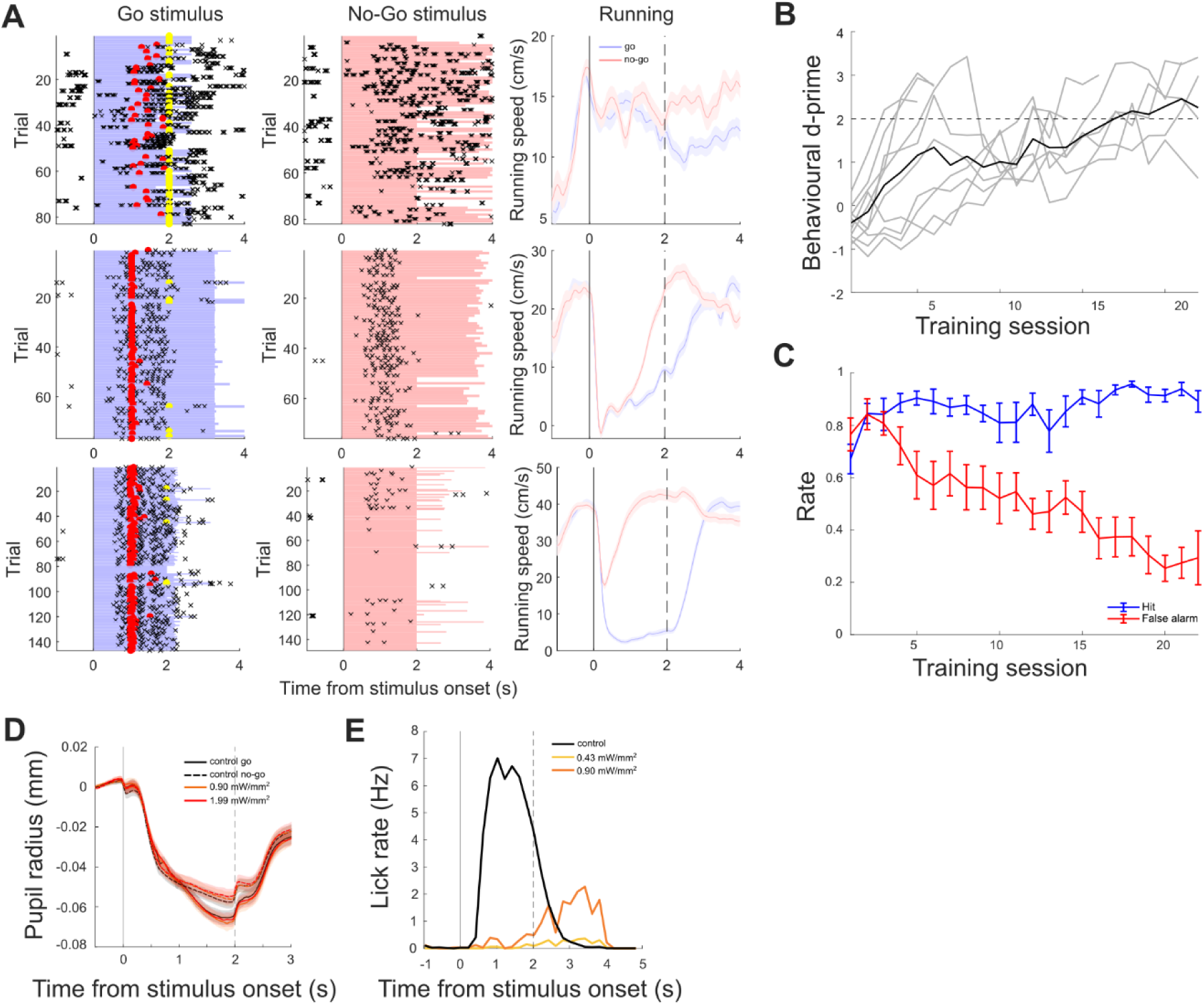
A) Changes in licking and running speed profile over training in an example mouse. Licks (black crosses) are aligned to stimulus onset in rewarded ‘go’ trials (blue shading, left panels) and unrewarded ‘no-go’ trials (red shading, middle panels). Red dots, reward delivery triggered by licks during the response window; yellow dots, reward delivery following auto-reward trigger. Average running speed (right panels) aligned to stimulus onset, for rewarded ‘go’ and unrewarded ‘no-go’ trials for the same example sessions. Shading, SEM. B) Average behavioural performance (*d’*, see Methods) and C) hit and false alarm rates across training sessions. Grey lines, individual mice; error bars, SEM. D) Average pupil size across different laser powers (including 0 mW/mm^2^) in ‘go’ (continuous line) and ‘no-go’ (interrupted line) conditions in WT mice. Shading, SEM. E) Lick rate during the presentation of stimulus only (black) or laser stimulation only (colour) in PV-ChR2 mice, indicating that mice cannot detect the laser stimulation alone in the absence of visual stimulus.

**Figure S2.**
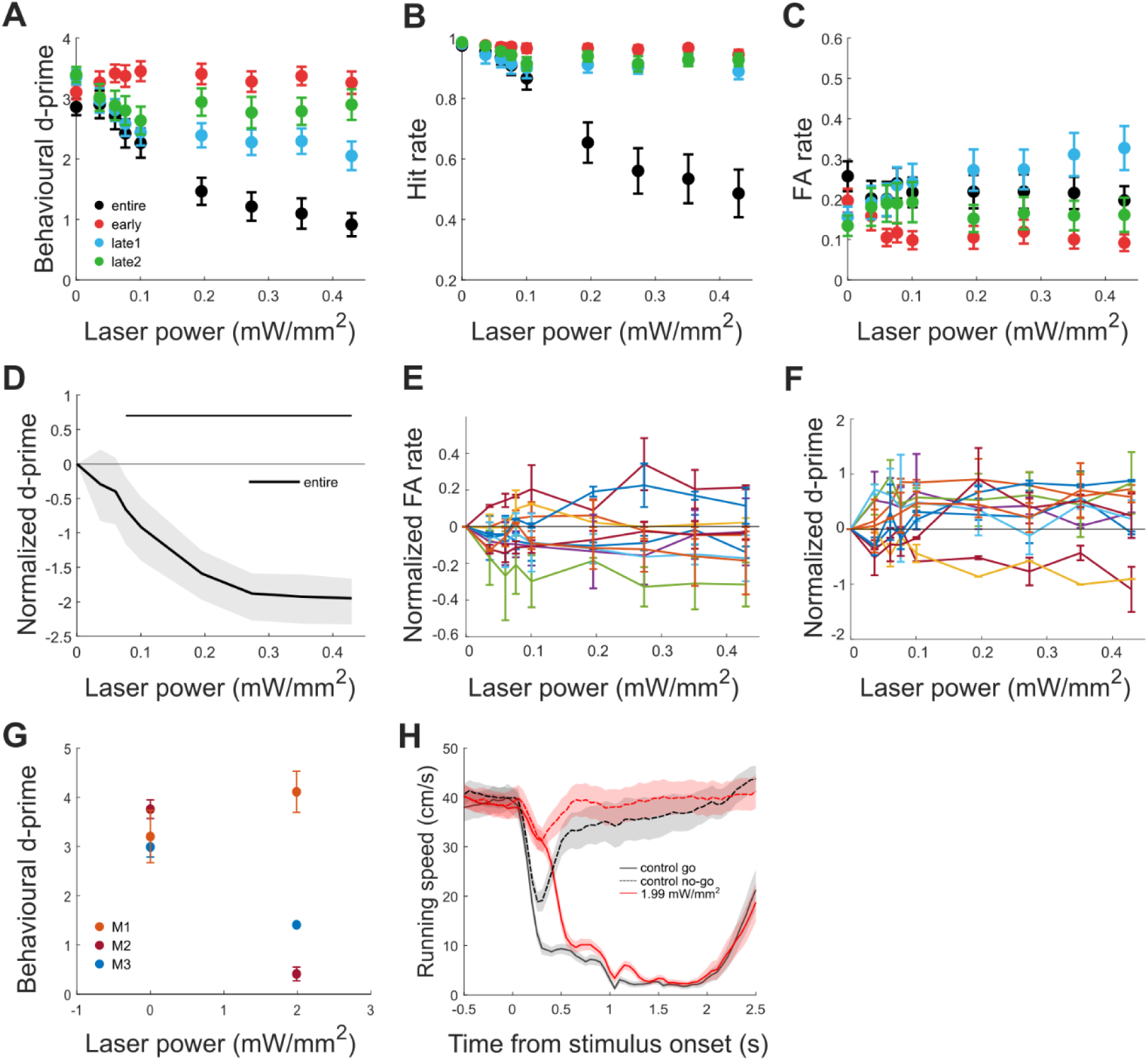
A) Average discrimination performance, *d’*, B) hit rate, and C) false alarm rate as a function of laser power for all time window conditions in the easy task. Error bars, SEM. D) Normalized average discrimination performance, *d’*, as a function of laser power for ‘entire’ window for sessions with lower hit rate (> 0.95). Shading, 95% CI of bootstrap distribution. E) Normalized average false alarm rate in the ‘entire’ window, and F) discrimination performance, *d’*, in the ‘early’ window in the easy task. Colour lines, individual mice (N = 10, 32 sessions); error bars, SEM. G) Average discrimination performance, *d’*, in three mice, where high laser power (2 mW/mm^2^) was used to silence the ‘early’ window (6 sessions). As expected, two of three mice showed impaired performance. Error bars, SEM. H) Average running speed for ‘go’ (continuous) and ‘no-go’ (interrupted) conditions for the mouse which retained high performance when the ‘early’ window was silenced at 2 mW/mm^2^. Running was delayed relative to stimulus presentation. Shading, SEM.

**Figure S3.**
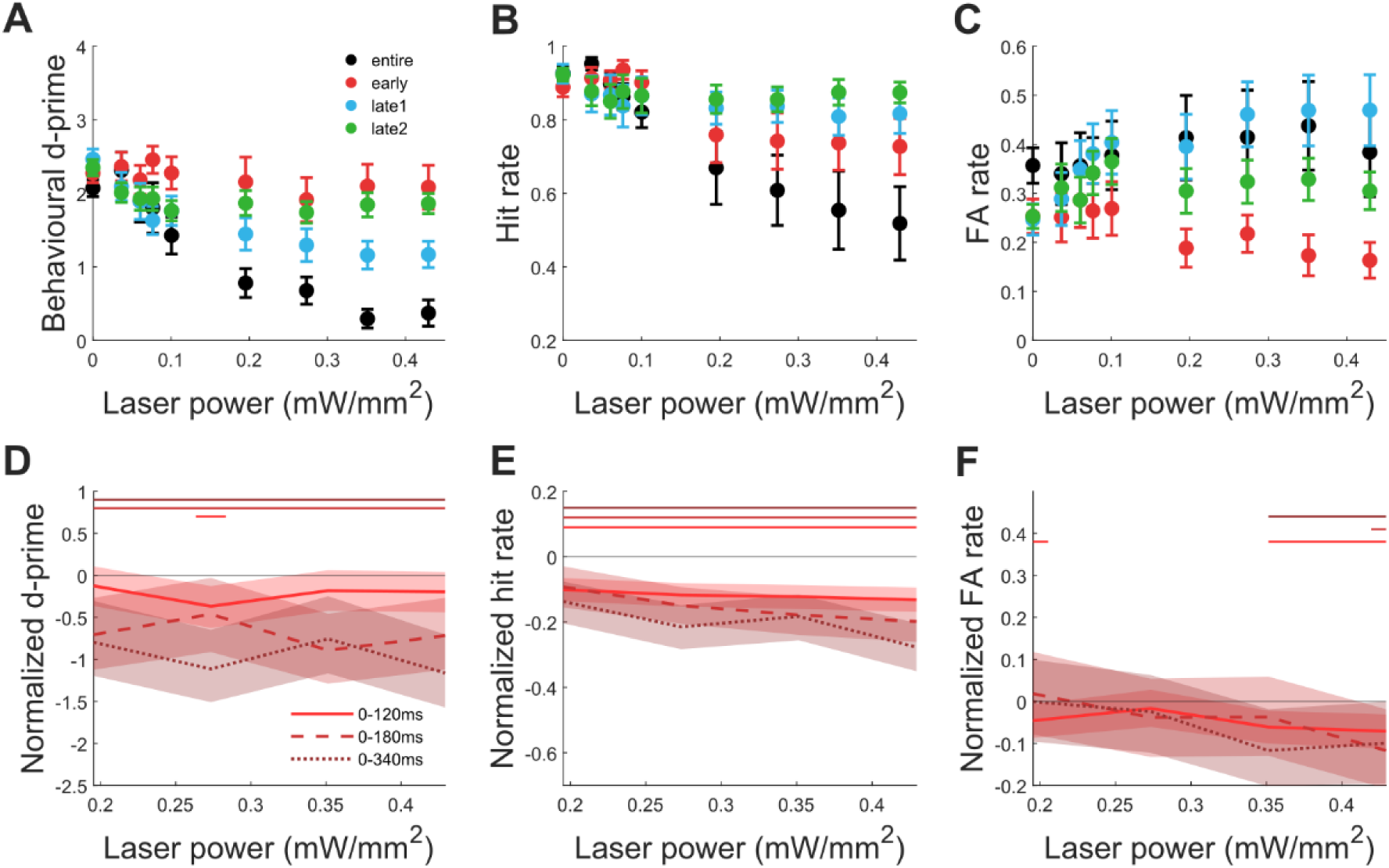
A) Average discrimination performance, *d’*, B) hit rate, and C) false alarm rate as a function of laser power for all time window conditions in the difficult task. Error bars, SEM. D) Normalized average discrimination performance, *d’*, E) hit rate, and F) false alarm rate as a function of laser power for the original (0 – 120 ms) and two new (0 – 180 ms and 0 – 340 ms) ‘early’ windows (N = 4 sessions). Shading, 95% CI of bootstrap distribution.

